# Two-photon microscopy *in vivo* reveals brain vessel type-specific loss of glycocalyx caused by apoM/S1P signaling impairment

**DOI:** 10.1101/2022.04.11.487803

**Authors:** Krzysztof Kucharz, Mette Mathiesen Janiurek, Christina Christoffersen, Martin Lauritzen

## Abstract

Increases in adsorptive mediated transcytosis (AMT) at the blood-brain barrier (BBB) are linked to many brain disorders. In a healthy brain, AMT is suppressed by sphingosine-1-phosphate (S1P) receptor 1 (S1PR1) signaling. Low levels of S1P lead to a rise in AMT, but the mechanisms are incompletely understood. Here, we explored whether the rises in AMT are caused by the loss of the endothelial glycocalyx (gcx). We used two-photon microscopy in mice with low S1P plasma levels (*Apom*^*-/-*^) and developed a novel photobleaching approach to measure gcx *in vivo* at distinct classes of cerebral microvessels, i.e., arterioles, capillaries and venules. We show that S1P signaling impairment reduced gcx in arterioles but not in other vessel segments. The location of gcx loss corresponded to the vascular topology of AMT increases. The S1PR1 agonist SEW2871 restores low levels of AMT in *Apom*^*-/-*^ mice but did not restore the gcx within the same time window. We propose that while the gcx loss may contribute to AMT increase, restoring gcx is not necessary for AMT to return to normal. These data establish a new imaging method to study gcx in the living mouse brain, demonstrate zonation of gcx in cerebral microvessels, and suggest differences in vascular susceptibility to gcx loss in disease states.

## INTRODUCTION

The blood-brain barrier (BBB) protects the fragile brain environment from toxins and pathogens circulating in the bloodstream. The BBB is composed of the brain endothelial cells (BECs) lining cerebral microvessels, non-cellular components, e.g., glycocalyx, and cells supporting BECs function: astrocytes and pericytes [1, 2]. In a healthy brain, the BBB permeability is low due to a tight seal of junctional complexes (JCs) between neighboring BECs [3], BECs efflux pumps [4], and ongoing suppression of vesicular transport across the BECs, i.e., transcytosis [5-7]. Transcytosis comes in two major modalities: the specific, receptor-mediated transcytosis (RMT) and the non-specific, adsorptive-mediated transcytosis (AMT) [5]. In disease states, AMT is no longer suppressed, causing a progressive build-up of toxic levels of albumin and other plasma constituents in the brain parenchyma before evident changes in the BBB structure. This has been evidenced in aging [8], stroke [9, 10] and may precede and facilitate neurodegeneration in chronic brain disorders [11-13].

We have recently shown that sphingosine-1-phosphate (S1P) receptor 1 (S1PR1) is a potent, homeostatic suppressor of AMT [7]. S1P is a signaling sphingolipid and the ligand for three main receptors at the brain BECs: S1PR1, S1PR2 and S1PR3 [14-17]. The majority of S1P in blood is bound to apolipoprotein-M (apoM) [18]. In this form, S1P activates preferentially S1PR1, but in the absence of apoM, the S1P levels are low, and S1PR1 signaling is impaired [17-19]. These conditions compromise the BBB, i.e., *Apom*^*-/-*^ mice exhibit both, increase in BBB paracellular permeability and excessive albumin transport to the brain via AMT [7, 20], but only in arterioles, while capillaries and venules remain unaffected. AMT in *Apom*^*-/-*^ mice returns to normal within 150 min in response to selective agonist of S1PR1 [7], but the mechanisms are unknown.

The S1P signaling is also involved in the maintenance of the glycocalyx (gcx) [21], a dense gel-like meshwork of glycoproteins and glycolipids that coats the luminal side of the BECs [22, 23]. The gcx acts as a mechanosensor [21, 24], signal transducer at BECs [21, 25-27], and a structural fence for blood-borne cell infiltration to the brain [28-30]. The gcx also constitutes a non-cellular component of the BBB [31], acting as a “molecular sieve” that hinders the diffusion of large, e.g., ∼30 kDa molecules across the BBB [32]. The negative charge of gcx enhances its structural barrier properties by repelling from BECs negatively-charged molecules in circulation, e.g., albumin, limiting their availability for AMT [33, 34].

Given that S1PR1 is expressed preferentially at arterioles [35], and arterioles exhibit both the highest gcx content in cerebral microvasculature [36] and susceptibility to AMT increase during impaired S1PR1 signaling [7], we hypothesized that the AMT increase could be explained by the loss of gcx, whereas return to normal AMT upon S1PR1 agonism was linked to restoration of gcx.

In order to examine the gcx, we used two-photon (2PM) imaging *in vivo*. The gcx is an elusive structure, typically lost during chemical tissue processing for immunohistochemistry (IHC) and transmission electron microscopy (TEM) [22, 23]. So far, the 2PM quantitative assessment of gcx *in vivo* has been either limited to a subset of capillaries [32] or perturbed by the background fluorescence of gcx labeling dyes, which circulate in an unbound form in the bloodstream [36]. Our first objective was to circumvent these issues. We developed a novel 2PM photobleaching approach to quantify the extent of gcx changes *in vivo* and at different categories of cerebral microvessels, i.e., arterioles, capillaries, and venules, irrespective of the background fluorescence of the unbound, blood-circulating dye. Our second objective was to determine: i) whether S1PR1 signaling impairment leads to the loss of gcx; ii) whether the loss of gcx occurs selectively at distinct types of cerebral microvessels, and if so, whether its topology corresponds to vascular locations of AMT increase; iii) and whether S1PR1 stimulation restores gcx, which might explain the return to normal levels of AMT in *Apom*^*-*/-^ mice [7].

We reveal that S1P signaling impairment in *Apom*^*-/-*^ mice leads to a degeneration of gcx in arterioles, which corresponded to the location of AMT increase [7]. However, stimulation of S1PR1 with the selective agonist SEW2871, which normalizes AMT within 150 min [7], did not reestablish the gcx within the same time frame. We conclude that although the loss of gcx may contribute to AMT increase in *Apom*^*-/-*^ mice, the reconstitution of the gcx is not critical for a low rate of AMT.

## MATERIALS & METHODS

### Animals

All experiments were performed in mice *in vivo*. The surgical and imaging procedures were approved by The Danish National Committee on Health Research Ethics and were performed according to the ARRIVE guidelines. To visualize gcx in relation to BECs *in vivo*, we used 23-26 g 16 weeks old Tg(TIE2GFP)287Sato/J transgenic reporter mice (Tie2-GFP; cat #003658, Jackson Laboratory)[37]. The experiments addressing apoM/S1PR1 signaling pathway and gcx were performed in 22-28 g 14-20 weeks old *Apom*^*-/-*^ mice [38], using C57Bl/6 (wild-type, WT) age- and gender-matched littermates as controls. The mice were genotyped to ascertain a double negative *Apom* allele as previously described [38]. All animals had free access to water and a chow diet and were housed in ventilated cages with a 12 h light / 12 h dark cycle. The animal facility had been accredited by the Association for Assessment and Accreditation of Laboratory Animal Care (AAALAC) and the Federation of Laboratory Animal Science Associations (FELASA).

### Preparative microsurgery

During all steps of surgery and imaging, the temperature of the animals was maintained at 37°C via a heating pad with feedback from a rectal thermal probe coupled to a homeothermic controller unit (EZ-TC-1000 Mouse Proportional). The animals were anesthetized with intraperitoneal (i.p.) injection of xylazine (10 mg/kg animal) and ketamine (60 mg/kg animal) with subsequent i.p, ketamine injections (30 mg/kg animal) in 20-25 min intervals. All mice underwent a tracheotomy for mechanical respiration (180-220 μl volume; 190-240 strokes/min; MiniVent Type 845 ventilator, Harvard Apparatus) to ascertain optimal blood oxygenation and for real-time monitoring of end-tidal exhaled CO_2_ (Capnograph Type 340, Harvard Apparatus) during the surgery and experiments. Next, the left femoral vein was cannulated to allow infusion of anesthetics during imaging, and the femoral artery was cannulated to monitor mean arterial blood pressure (MABP; Pressure Monitor BP-1, World Precision Instruments) and for blood gasses sampling and delivery of fluorescent probes. Following that, all wounds were surgically stapled, and the animal was placed in a prone position (Fig. 1a).

**Fig.1.**
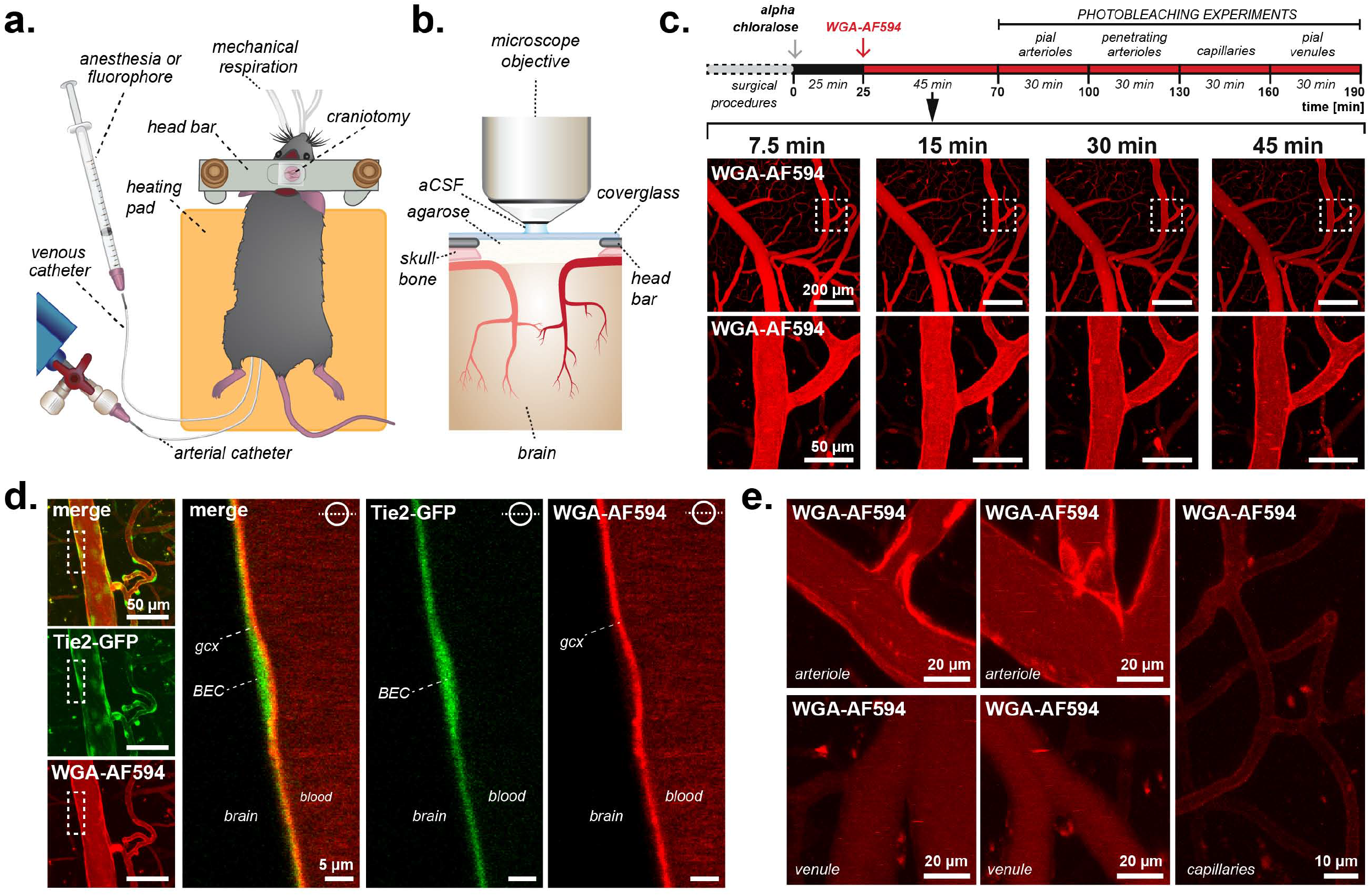
Gcx labeling in vivo. **a)** A mouse after the preparative microsurgery for 2PM imaging; **b)** Features of the cranial window; **c)** Experimental timeline (upper panel) and a time-lapse of the gcx labeling by WGA-AF594 (lower panel). The time above the images is relative to the WGA-AF594 injection. **d)** Spatial relation between endothelium (green) and gcx labeling (red). The BECs were delineated by GFP signal expressed in BECs in transgenic mice (Tie2-GFP). The WGA-AF594 signal was located at the luminal side of BECs, at the expected location of gcx. **e)** Enriched presence of gcx was present at the branching points of arterioles but not at venules or capillaries. **All panels:** *gcx*= glycocalyx; *BEC*=brain endothelial cell; all images are maximum-intensity projected Z-stacks unless indicated by a circular inset that denotes the position of the imaging plane in relation to the vessel cross-section. Panels a) and b) were reused from *Janiurek et al. 2019* under CC-BY 4.0 license.

### Cranial window surgery

To allow the two-photon imaging, a craniotomy was performed. First, the scalp was removed, and the periosteum of the exposed skull was gently peeled off using a cotton bud pre-soaked in FeCl_3_ (0.62 M). Next, the skull was dried and glued (Loctite Adhesives) to a custom-made metal plate that was subsequently mounted onto the imaging stage insert. The craniotomy (Ø = 4 mm) was performed using a diamond dental drill (7000 rpm) at the right parietal bone, at the location of the somatosensory cortex (3 mm lateral to the sagittal suture, 0.5 mm posterior to the coronal suture). To prevent heat damage during the drilling process, the skull surface was repeatedly cooled using room-temperature saline. Next, the bone flap was lifted, and the exposed brain surface was gently rinsed with artificial cerebrospinal fluid (aCSF; in mM: NaCl 120, KCl 2.8, Na_2_HPO_4_ 1, MgCl_2_ 0.876, NaHCO_3_ 22, CaCl_2_ 1.45, glucose 2.55, at 37°C, aerated with 95% air/5% CO_2_ to pH 7.4). The dura was carefully removed, and a drop of low-melting agarose (0.75% agarose, type III-A, Sigma-Aldrich) was applied to cover the brain surface. Lastly, a glass coverslip (Menzel-Gläser, 24×66mm, #1.5) was positioned onto the skull opening to allow optimal imaging and seal the craniotomy (Fig. 1b).

Following the microsurgery, the imaging insert with the animal was moved to the microscope stage, and the anesthesia was switched to alpha-chloralose (Sigma-Aldrich) administered by continuous infusion via a venous catheter (50 mg/hour/g animal). The change from ketamine to alpha-chloralose was to minimize the anesthesia effect on NMDAR-dependent neurotransmission in the brain. The animal was allowed to rest for 25 minutes before the imaging.

### Monitoring physiological parameters

The percentage of exhaled CO_2_ was monitored in real-time (Capnograph Type 340, Harvard Apparatus), and the respiration volume and stroke frequency were adjusted when necessary. Prior to imaging experiments, the animal’s physiological state was further evaluated by sampling 50 μl of blood from the femoral artery for O_2_, CO_2_ blood gasses and blood pH measurements (ABL 800Series, Radiometer). The range of parameters considered to be physiological was: 95–110 mmHg for pO_2_, 35 – 40 mmHg for pCO_2_, 7.35 – 7.45 for pH (all blood sample measurements); and 50 – 80 mmHg for MABP, 2.2 – 2.8% for exhaled CO_2_ (all real-time monitoring), based on our previous two-photon *in vivo* assessments of neurovascular coupling [39], neuronal activity [40] and the BBB [41].

### Imaging setup

The two-photon (2PM) *in vivo* imaging was performed using Leica SP5 upright laser scanning microscope equipped with 20 × 1.0 NA water-immersion objective (all Leica Microsystems) and MaiTai Ti:Sapphire laser (Spectra-Physics). The emitted light was split by TRITC/FITC dichroic mirror, and collected after 525–560 nm and 560–625 nm bandpass filters, for second harmonics/GFP and Alexa Fluor™ 594 imaging channels, respectively. All data was acquired as multi-channel 16-bit images using the LAS AF imaging software suite (v. 3.5; Leica Microsystems).

### Glycocalyx labeling

The glycocalyx was labeled *in vivo* by a single bolus intraarterial (i.a.) injection of wheat germ agglutinin (WGA) -Alexa Fluor^™^ 594 conjugate (0.004 mg/g BW of 1% WGA-Alexa Fluor™ 594, Invitrogen) via the femoral artery catheter and imaged with the laser tuned to 850 nm. The time-lapse overview images were collected as the Z-stacks from the cortical volume 775 μm × 775 μm × 100 μm with 2.5 μm Z-step (2048 × 2048 pixel resolution), then exported as multi-layer 16-bit tiff files to ImageJ (v. 1.52, NIH), and presented as maximum intensity projections along the Z-axis. The optimal time for gcx imaging was at 45 minutes post WGA-Alexa Fluor 594 injection, based on the vessel lumen-to-gcx signal ratio in arterioles (Fig. 1c). In the overview images showing endothelium (Tie2-GFP mice), the GFP signal was acquired at 850 nm simultaneously with WGA-Alexa Fluor 594.

### Glycocalyx photobleaching

To quantify the gcx labeling, we developed a protocol based on photobleaching of the gcx-bound WGA-Alexa Fluor 594 (Fig. 2a). The purpose of photobleaching was to differentiate the background fluorescence of the brain parenchyma and the blood-circulating dye from the fluorescence of labeled gcx. The photobleaching protocol consisted of three steps. First, we acquired a *pre-bleaching* image from a small vessel area (129 μm × 129 μm; 512 × 512 pixels) that encompassed gcx at the blood-brain interface at the focal plane located at the broadest section of the vessel. The fluorescence at the gcx location contained the fluorescence from WGA-Alexa Fluor 594-labeled glycocalyx, with the addition of background fluorescence of circulating (gcx-unbound) dye from the vessel lumen (on the luminal side of gcx), and the brain parenchyma autofluorescence (on the abluminal side of gcx). Next, we performed rapid photobleaching of a small region of interest (bROI; 15 μm × 15 μm; 512 × 512 pixels) at the blood-brain interface by fast bi-directional scanning of the bROI for consecutive 12 frames at 0.375 s intervals (laser power at the sample = 35 mWatt at 850 nm)(Fig. 2a, b). Immediately after, we collected a *post-bleaching* image from the same vessel area using the same parameters as for the *pre-bleaching* image. The *post-bleaching* image contained fluorescence attributed exclusively to background fluorescence and blood-circulating WGA-Alexa Fluor 594 (Fig. 2a,c).

**Fig. 2.**
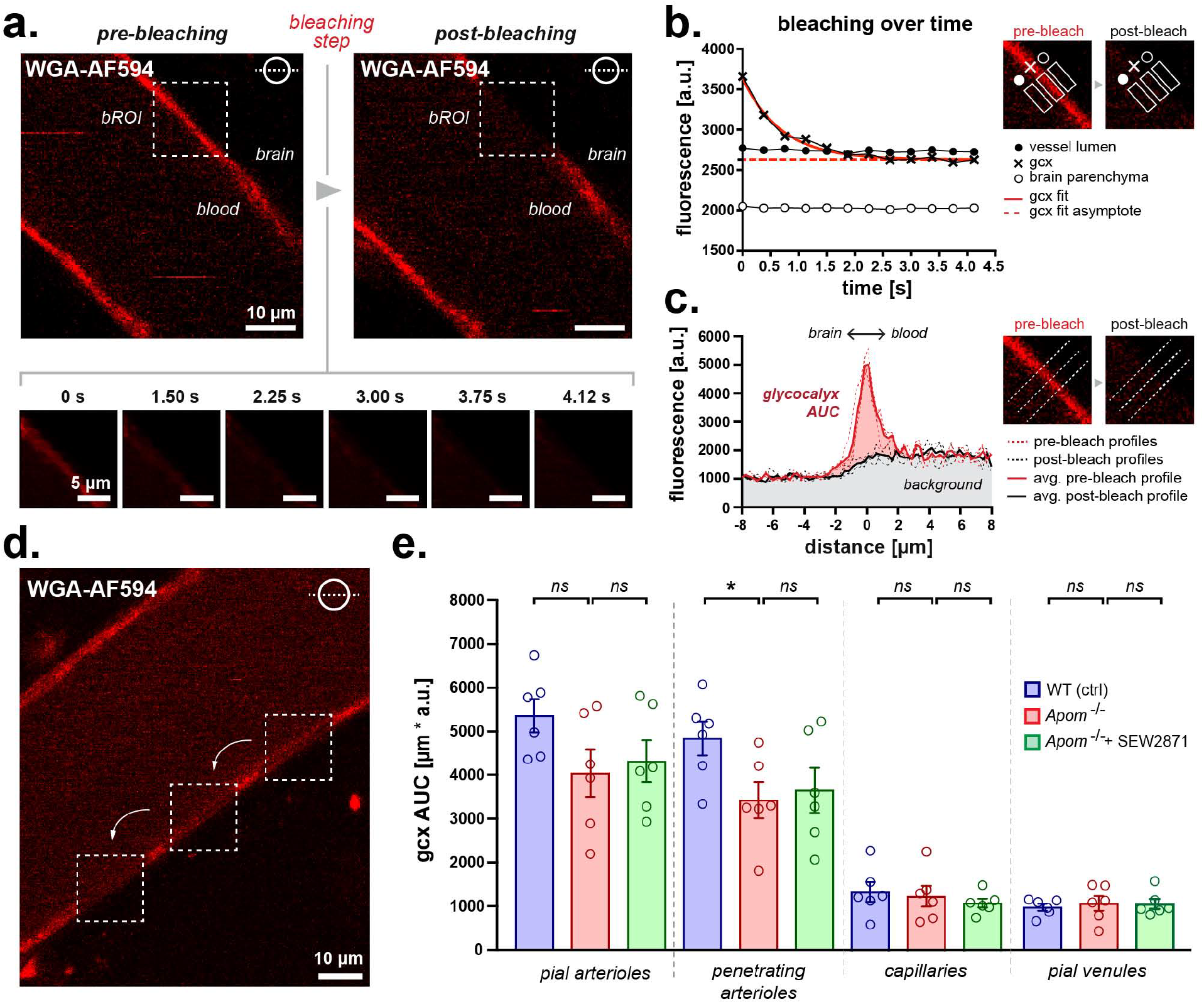
2PM photobleaching reveals vessel-specific loss of gcx in *Apom*^*-/-*^ mice, with no rescue effect by S1PR1 stimulation. **a)** The principle of photobleaching protocol. A square bleaching region of interest (bROI) is selected at the location of gcx (pre-bleaching image). Next, the bROI is bleached during 4.12 s of a continuous scan (lower panel), resulting in fluorescence loss of the gcx within the area (post-bleaching image). **b)** During the photobleaching step, the fluorescence from the gcx rapidly approached the asymptote (red dashed line), indicating efficient quenching. The bleaching preserved the background signal of circulating WGA-AF594 measured from neighboring vessel lumen (black dots) and brain parenchyma (white circle). **c)** The AUC of the gcx signal (red) was calculated from the difference between the average pre-bleach (red trace) and average post-bleach (black trace) fluorescence intensity profiles obtained from three profile measurements before and after the bleaching, respectively. The image insets show the orientation of fluorescence intensity profiles at the vessel wall. **d)** The photobleaching protocol was repeated sequentially along the vessel wall. **e)** Comparisons between gcx AUC between WT, *Apom*^*-/-*^, and SEW2871-treated *Apom*^*-/-*^ mice. The S1PR1 signal deficiency led to a decrease in gcx in penetrating arterioles, which was not rescued by S1PR1 agonism. Other vessel segments were unaffected by S1PR1 signaling impairment or SEW2871 treatment. Each symbol denotes a single mouse (n=6/group). Data are presented as mean with SEM. ns= non-significant, with asterisk denoting *p<0.05 (Mann-Whitney). **All panels:** *gcx*= glycocalyx; *bROI*=bleaching region of interest; *AUC*= area under the curve, *a*.*u*.=arbitrary fluorescence unit. The circular inset denotes the position of the imaging plane in relation to the vessel cross-section.

Next, the imaging data was exported to ImageJ (v. 1.52, NIH) as multi-layered 16-bit tiff files for further analysis. For each *pre-bleaching* image, we extracted three fluorescence intensity line profiles oriented perpendicular to the vessel wall. The first profile was located at the center of bROI, and the remaining two were offsetted from the center by -2.5 and 2.5 μm, respectively. The three profiles were averaged to obtain the representative *pre-bleaching* profile for a single bROI. The same procedure was then repeated for the *post-bleaching* image to obtain a representative *post-bleaching* profile for the same bROI (Fig. 2c). The fluorescence signal is additive; therefore, we next subtracted *post-* from the *pre-bleaching* representative profiles to obtain a signal originating exclusively from gcx. Given the diffraction limit of 2PM, instead of estimating gcx thickness, we provided a single measure of gcx that convolved both gcx density and length. The amount of gcx was expressed as the area under the curve (AUC) of a single peak at the expected location of gcx (Fig. 2c) and calculated using the trapezoid method (GraphPad Prism, v. 8).

The photobleaching protocol was performed for multiple locations, i.e., multiple bROIs, within each vessel, progressing along a vessel wall (Fig. 2d). For each animal, the data were first averaged for multiple bROIs within the analyzed vessel and then between the same vessel types.

### S1PR1 stimulation with SEW2871

To stimulate S1PR1, we used the specific agonist, SEW2871 [42]. To better relate our results to our previous findings on AMT [7], the animals were imaged 150 min after a single i.p. bolus injection of SEW2871 (10 μg/g BW). SEW2871 was dissolved to 10 mg/ml in DMSO and diluted to 1 mg/ml in PBS before injection. The vehicle solution was PBS containing 10% DMSO.

### Syndecan-1 blood plasma measurements

The 40 μl of blood was collected from the tail into the K2-EDTA containing microcuvettes (Starstedt) and immediately centrifuged at 2000×g for 10 minutes to isolate plasma. Next, the plasma was transferred into the fresh microcuvette and stored at -80°C for further use. The concentrations of blood plasma syndecan-1 were measured using a Mouse SDC1 ELISA Kit (#LS-F28946, LifeSpan BioSciences, Inc.) according to the manufacturer’s instructions, using a PowerWave XS microplate spectrophotometer (BioTek Instruments Inc.). The assessment from all timepoints from each mouse was performed in a single run, with each timepoint measured in duplicates.

### Statistical analysis

The number of mice was considered as the number of independent replicates. The 2PM datasets were tested for normal distribution with the D’Agostino-Pearson method (GraphPad Prism, v. 8). Limited sample size prevented the unequivocal determination of normality; therefore, we used two-tailed Mann–Whitney rank-sum tests, with Bonferroni post-hoc corrections for multiple comparisons between the groups. To determine the effects of genotype or treatment on syndecan-1 levels in the blood, we used two-way ANOVA, with Bonferroni post-hoc correction for multiple comparisons. The experiments were performed in a randomized manner by an observer not blinded to experimental conditions.

### Exclusion criteria

All animals exhibiting low blood pressure (<50 mmHg) or substantial brain movement during the imaging were excluded from the study (2 out of 21 animals total). No animals were excluded from ELISA experiments (n= 18 animals total). We observed no attrition due to gcx labeling, photobleaching procedures, or SEW2871 treatment.

## RESULTS

### Glycocalyx labeling *in vivo*

Our first objective was to optimize the gcx labeling for 2PM *in vivo*. The brain microvasculature was imaged via a cranial window at the location of the somatosensory (i.e., barrel) cortex (Fig. 1a, b). Following the microsurgery, we injected WT mice with wheat germ agglutinin conjugated to Alexa Fluor™ 594 (WGA-AF594) via a previously mounted arterial catheter. The blood-circulating WGA-AF594 did not instantaneously label the gcx; instead, we observed a gradual fluorescence increase at the blood-brain interface (Fig. 1c). The signal from gcx-bound WGA-AF594 could be clearly distinguished from gcx-unbound dye in plasma at 45 min post-injection. Given that the labeling of gcx is time-sensitive, the imaging was performed in a time-consistent manner. Specifically, the pial arterioles were always assessed first, at 45 min post-injection and for 30 min, then penetrating arterioles, capillaries, and venules in consecutive 30 min periods (Fig. 1c). In addition, we ascertained that the WGA-AF594 fluorescence originated from the expected location of gcx, i.e., the luminal side of BECs, using Tie2-GFP mice expressing the green fluorescent protein (GFP) in the brain endothelium (Fig. 1d). Our first observation was that the gcx presence was more pronounced in arterioles compared to venules, indicating a heterogeneous distribution of gcx in the vascular tree. However, the gcx coverage could be non-uniform within an individual vessel. In particular, we discovered enriched gcx labeling at the branching points of larger (Ø 20-40 μm) arterioles. Notably, this pattern was never present at diameter-corresponding branches of coalescing venules (Fig. 1e).

### Glycocalyx photobleaching

Next, we devised a method to quantitatively characterize the gcx at the BBB. Fluorescence measurements of WGA conjugates is the backbone of quantitative assessments of gcx *in vivo* [36, 43]. However, the signal collected at the location of gcx consists not only of the fluorescence from gcx-bound WGA conjugates, but also contains a significant contribution from other sources, e.g., collagen-rich vessel walls and the unbound dye circulating in the bloodstream. This perturbs gcx measurements and may lead to an overestimation of gcx expression, especially for vessels with low gcx content and high background signal. Here, we used photobleaching to minimize this interference and extracted the fluorescence signal attributed exclusively to WGA-AF594 -labeled gcx. We fast-scanned small areas at the vessel boundaries, which resulted in photobleaching of the WGA-AF594 bound to gcx (see Methods)(Fig. 2a), but without significant effects on the background fluorescence, i.e., fluorescence measured from unbound WGA-AF594 in the bloodstream and autofluorescence from brain parenchyma (Fig. 2a, b). The photobleaching caused WGA-A594 fluorescence at the BBB to rapidly decline towards its asymptote, indicating fast and effective quenching of gcx-bound dye, but did not compromise the BBB integrity, as ascertained by the lack of extravasation of WGA-AF594 from the blood to the brain parenchyma (Fig. 2b). The difference between *pre-* and *post-bleaching* fluorescence intensity profiles was used to extract the signal corresponding to gcx-bound WGA-AF594 (see Methods)(Fig 2c). Given that the gcx thickness can approach the diffraction limit of the 2PM (0.42 μm for 850 nm **λ**_excitation_ and 1 NA objective), we refrained from estimating the gcx thickness. Instead, as the measure of gcx, we used the area under the curve of the gcx bleached signal from fluorescence profile plots (gcx AUC, red)(see Methods)(Fig. 2c), which is proportional to both the thickness and the density of the gcx. The procedure was repeated sequentially along the vessel wall for each vessel to obtain information on the average gcx coverage for a particular vessel (Fig 2d).

### S1PR1 signaling deficits induce loss of gcx, but only in arterioles

Our second goal was to determine whether S1P signaling impairment affects the gcx at the BBB. In line with our previous study [7], we used *Apom*^*-/-*^ mice that exhibit a reduction of S1P levels in plasma [38]. Ongoing S1PR1 receptor activity prevents the gcx shedding *in vitro* [21], but cultured cells do not recapitulate the functional and structural zonation of the BBB. Therefore, we analyzed the gcx across pial and penetrating arterioles, capillaries and pial venules, and up to 90 μm below the pial surface. Using the photobleaching approach, we found that impaired S1P signaling decreased gcx in *Apom*^*-/-*^ mice compared to WT controls (Fig 2e). This was observed in penetrating arterioles (3723±410.9 vs. 4842±387.9 μm*a.u., respectively, p=0.0152, n=6 mice/group, Mann-Whitney), with a strong trend for gcx reduction in pial arterioles (4046±550.0 vs. 5365±380.9 μm*a.u., p=0.0649, n=6 mice/group, Mann-Whitney). However, in contrast to penetrating arterioles, S1P signaling impairment had no effect on the gcx coverage in capillaries (1329±169.8 vs. 1566±251.6 μm*a.u., p=0.8182, n=6 mice/group, Mann-Whitney) and venules (1075±166.4 vs. 989±77.69 μm*a.u., p=0.8182, n=6 mice/group, Mann-Whitney). These results indicate that S1P signaling impairment is accompanied by the loss of gcx in penetrating arterioles but not in other microvascular segments.

### S1PR1 agonism does not restore the gcx on a short time scale

Next, we tested whether S1PR1 agonism reinstates the gcx. We have previously shown that S1PR1 selective agonism rapidly (<150 min) restores the low rate of AMT in *Apom*^*-/-*^ mice [7]. Given that gcx is a barrier for large molecules [31, 32], restoring gcx within the same time frame might explain the return to normal AMT. We used the same treatment regime that normalizes AMT, i.e., single bolus i.p. injection with SEW2871 (10 μg/g BW)[7], and imaged gcx in *Apom*^*-/-*^ mice 150 min thereafter.

Our 2PM photobleaching results showed that the SEW2871 did not restore the gcx (Fig 2e). There was no significant difference in gcx in penetrating arterioles between treated and non-treated *Apom*^*-/-*^ mice (4320±485.1 vs. 4046±550.9 μm*a.u., respectively, p=0.5887, n=6 mice/group, Mann-Whitney). Noteworthy, capillaries and venules that were unaffected by S1PR1 signaling deficit also did not respond to the treatment, e.g., with an increase in the gcx content (pial arterioles: 4320±485.1 vs. 4046±550.9 μm*a.u., p=0.5887; capillaries 1253±103.0 vs. 1329±169.8 μm*a.u., p=0.8182; and pial venules 1063±112.7 vs. 1075±166.4 μm*a.u., p>0.9999; for treated-vs. non-treated *Apom*^*-/-*^, respectively, n=6 mice/group; all Mann-Whitney)(Fig 2e).

WGA-conjugated fluorescent dyes are suitable for labeling the main constituents of the gcx filament structure, i.e., glycosaminoglycans (GAGs) [25, 44], but they do not visualize BEC membrane proteins that anchor GAGs to the BECs. Therefore, complementarily to 2PM, we measured blood plasma levels of syndecan-1, a transmembrane protein component of the gcx [23, 45, 46]. The WT and *Apom*^*-/-*^ mice were assessed longitudinally, i.e., in 60 min intervals and up to 240 min after a single bolus i.p. injection with vehicle or SEW2871 (10 μg/g BW).

First, we found that the *Apom*^*-/-*^ mice exhibited a substantial reduction in circulating syndecan-1 compared to WT littermates (WT vs. *Apom*^*-/-*^; ^###^p<0.001; n=4 vs. 5 mice, respectively; two-way ANOVA; F(1,7)=43.37)(Fig 3a). Increased concentrations of syndecan ectodomain in blood plasma are typically interpreted as signs of the acute shedding of gcx. Given that the S1P facilitates the gcx synthesis [47], our results indicate chronic low levels of gcx components in *Apom*^*-/-*^ mice rather than decreased shedding compared to WT controls.

**Fig. 3.**
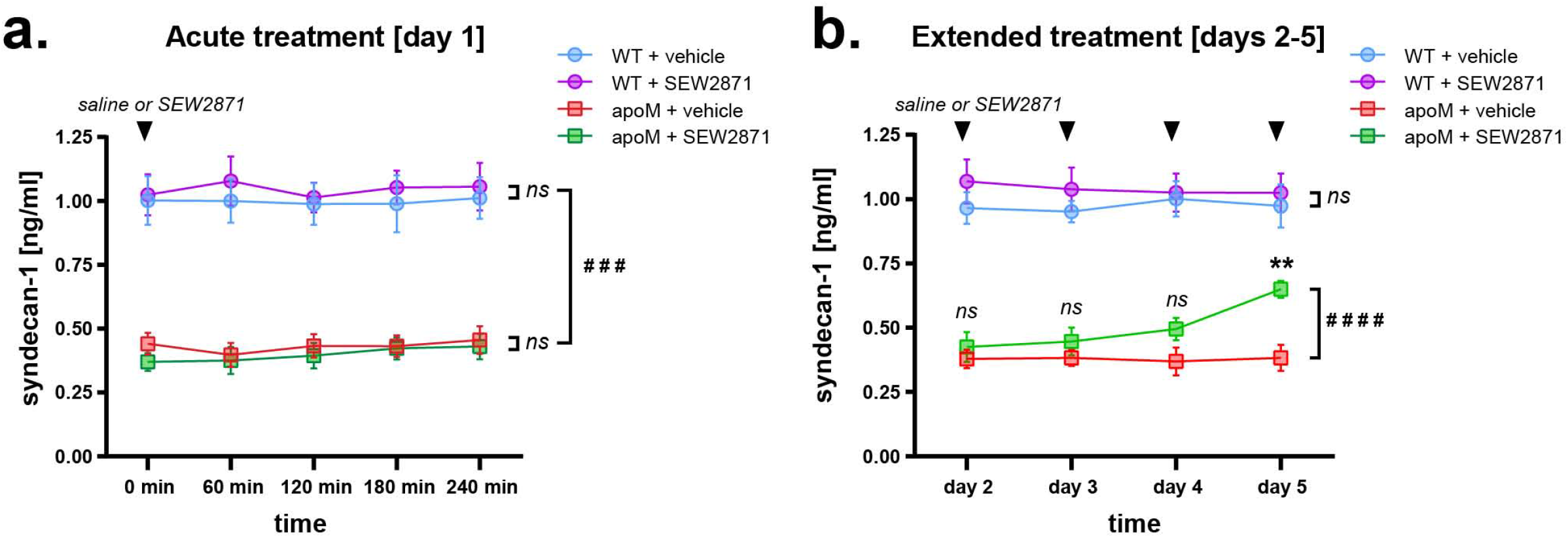
Longitudinal assessment of syndecan-1 plasma levels during S1PR1 agonism in *Apom*^*-/-*^ mice. **a)** The plasma concentrations of circulating syndecan-1 were significantly decreased in *Apom*^*-/-*^ compared to WT mice, and the S1PR1 agonism did not affect syndecan-1 levels within the time corresponding to 2PM experiments (up to 240 min). **b)** Recurrent treatment with SEW2871 triggered a delayed increase in circulating syndecan-1 in *Apom*^*-/-*^ compared to non-treated *Apom*^*-/-*^ mice, and did not affect WT mice. **All panels:** data are averages with SEM; underlying numbers denote significance values of multiple comparisons. ^####^*p*<0.0001; ^###^*p*<0.001 (two-way ANOVA); \*\**p*<0.01 (two-way ANOVA multiple comparisons with Bonferroni post-hoc correction); *ns*= non-significant. n_WT_=4; n_WT+SEW2871_=4; n_Apom-/-_=5; n_Apom-/-+SEW2871_=5 mice.

Next, we tested whether S1PR1 agonism affected syndecan-1 levels in *Apom*^*-/-*^ mice in a time window corresponding to AMT return to normal [7]. We found no effect of SEW2871 on gcx in both *Apom*^*-/-*^ (*Apom*^*-/-*^ vs. *Apom*^*-/-*^+SEW2871; p=0.4975; n=5 mice/group; two-way ANOVA, F(4,32)=0.8614), and WT mice (WT vs. WT+SEW2871; p=0.9421; n=4 mice/group; two-way ANOVA, F(4,24)=0.1884)(Fig. 3a). In line with 2PM photobleaching data, this result indicated the lack of gcx restoration upon selective S1PR1 agonism in *Apom*^*-/-*^ mice.

We extended the treatment for additional 4 days, with blood samples taken at 240 min after daily i.p. injections with vehicle or SEW2871 (10 μg/g BW) (Fig 3b). The S1PR1 agonism eventually increased syndecan-1 plasma levels (*Apom*^*-/-*^ vs. *Apom*^*-/-*^+SEW2871; ^####^p<0.0001; n=5 mice/group; two-way ANOVA, F(3,24)=14.39), but the changes were evident only at day 5 of the treatment (*Apom*^*-/-*^ vs. *Apom*^*-/-*^+SEW2871; ^**^p<0.01; n=5 mice/group; two-way ANOVA multiple comparisons with Bonferroni correction).

Taken together, our results indicate that S1P signaling deficiency induces selective loss of gcx in cerebral arterioles, corresponding to the location of AMT increase [7]. However, S1PR1 agonism that restores normal AMT within 150 min did not restore gcx within the same time frame, questioning the role of gcx as the bidirectional modulator of AMT.

## DISCUSSION

Impaired S1P signaling, disturbances in gcx, and detrimental increases in AMT are co-existing features of many brain disorders linked to BECs dysfunction. Our current, limited understanding of the mechanisms of these processes comes primarily from *in vitro* models of the BBB, which do not recapitulate the structural heterogeneity of the brain microvasculature and functional zonation of transport mechanisms across the BBB [7, 8, 41, 48]. In turn, assessments of gcx *in vivo* face a methodological challenge due to limitations of imaging techniques.

Here, we developed a 2PM fluorescence photobleaching approach to quantitatively characterize the gcx at different categories of cerebral microvessels. Using *Apom*^*-/-*^ mice, we show that S1P signaling impairment leads to a chronic decrease in gcx content at the BBB. Gcx reduction was present selectively at arterioles, which mirrored the vascular location of AMT increase in *Apom*^*-/-*^ mice evidenced in our previous study [7]. However, S1PR1 stimulation that rapidly, i.e., within 150 min, restores the physiological rate of AMT [7] did not reinstate the gcx within the same time frame. We suggest that although the loss of gcx is spatially correlated with increased AMT, the presence of gcx is not necessary to normalize transcytosis.

The extent of gcx coverage in the brain depended on vessel type rather than the vessel lumen diameter. The gcx distribution was non-uniform along the vascular tree, with the highest presence at arterioles, consistent with earlier reports [36]. In addition, we detected “hot-spots” of fluorescence from WGA-A594 at the branching points of arterioles, but this pattern was absent in size-corresponding venules. The blood pressure and blood cell velocity are highest in arterioles, subjecting arteriolar branching points to greater shear stress compared to corresponding locations at capillaries and venules [49, 50]. Recently, laminar shear stress has been found to stimulate local biosynthesis of gcx [51]. Given that the gcx facilitates efficient red blood cell flow within the microvessel lumen, the enriched presence of gcx at the arteriolar branching points may be an adaptive response, supporting optimal perfusion to downstream capillaries [52].

Next, using the 2PM photobleaching approach, we found that *Apom*^*-/-*^ mice exhibit a reduction in gcx. This result is in accord with previous studies showing that S1P facilitates the synthesis and inhibits shedding of the gcx *ex vivo* [47, 53], and that *Apom*^*-/-*^ mice exhibit low S1P plasma levels [17-19]. However, the overall dependency of gcx on S1P does not explain why only arterioles were susceptible to the gcx loss. One explanation is that rather than being linked with a general decrease of S1P in *Apom*^*-/-*^ mice [18], the reduction in gcx is linked with the hypostimulation of a particular S1P receptor, i.e., S1PR1. First, the reduction in apoM is thought to shift the signaling equilibrium from S1PR1 to S1PR2 and S1PR3, which may act antagonistically to S1PR1 [19]. Second, the transcriptomic data show that S1PR1 is expressed preferentially in arterioles rather than in capillaries and venules [35, 54]. The high expression of S1PR1 in arterioles may suggest that the gcx in this vessel segment is particularly dependent on preserved S1PR1 activity, but further evidence is needed to support this notion.

The gcx is a “structure in flux” determined by a balance between synthesis and shedding [22, 23]. Increased shedding of gcx accompanies aging [55, 56] and pathologies affecting vascular function, e.g., stroke [10, 57], sepsis [58, 59] and diabetes [60]. These conditions have a common denominator, impaired S1P signaling [61-65] and all feature an increase in AMT [8, 9, 11-13, 66]. The joint occurrence of S1P signaling deficits, gcx loss and AMT increase may suggest a causal relationship, but these features have never been cross-examined in the same experimental setting. We have previously shown that impairment of S1P signaling compromises the BBB. Specifically, the *Apom*^*-/-*^ mice exhibited an abnormal increase in AMT, and the effect was present only in arterioles [7]. Here, we discovered that the gcx decrease in *Apom*^*-/-*^ mice is also selective to arterioles, mirroring the vascular topology of AMT increase. The gcx is an intrinsic part of the BBB [31, 32]. Its dense meshwork of glycosaminoglycans (GAGs) constitutes a diffusional barrier for large, e.g., >30 kDa molecules [31, 32], and the anionic GAG components enhance barrier function by repelling negatively charged molecules from the luminal side of BECs [22, 23, 34]. The quintessential protein undergoing AMT is serum albumin [1, 5]. In a healthy brain, interactions of albumin with BECs are effectively hindered by gcx due to both albumin size (∼65kDa) and a negative charge at physiological pH (7.4) [33, 34][67]. Therefore, S1P signaling deficiency that causes a reduction in gcx may increase the probability of albumin interactions with BECs and the rate of endocytosis, which may, in part, explain the underlying mechanism of AMT disinhibition. S1P binds mainly to apoM, but albumin may carry up to 30 % of S1P in plasma. *Apom*^*-/-*^ mice have a level of S1P-albumin in plasma comparable to WT mice [18]. This suggests that the presence of S1P-albumin in *Apom*^*-/-*^ mice may not be enough to reduce AMT, but that this process may be an apoM-S1P specific mechanism.

Given that S1P signaling facilitates the synthesis of gcx [47], we explored whether S1PR1 agonism reinstates the gcx, which could underlie the rescue of the BBB function observed in our previous study [7]. We have previously shown that selective stimulation of S1PR1 with SEW2871 restores low, physiological levels of AMT in *Apom*^*-/-*^ mice in less than 150 min [7]. Here, we imaged the gcx in *Apom*^*-/-*^ mice subjected to the same treatment, but observed no recovery of gcx within the corresponding time. This indicated that while loss of gcx may facilitate AMT increase, reinstating the gcx structure is not a prerequisite to normalizing AMT, and other, indirect mechanisms may be in play.

The gcx is not only a passive barrier at the BBB but acts as a signal transduction component that can influence BECs function [21, 26, 27]. Syndecan-1 is a membrane proteoglycan and the backbone of the gcx structure that anchors GAGs to endothelium [23, 45, 46]. *In vitro* assays show that upon shedding, the remaining cytosolic fragment of syndecan-1 promotes Src-mediated phosphorylation of caveolin-1, and endocytosis [68-70]. Extrapolating to *in vivo*, this mechanism has been proposed to drive the fluctuations in transcytosis after stroke, where AMT increases simultaneously with gcx shedding and decreases synchronously with the restoration of syndecan-1 [10]. However, our results do not support syndecan-1 involvement in the recovery to physiological levels of AMT in *Apom*^*-/-*^ mice. We found persistently low levels of plasma syndecan-1 in *Apom*^*-/-*^ mice. This suggests a chronic reduction in the syndecan-1 total pool and, therefore, in the amount of shed syndecan-1 in *Apom*^*-/-*^ mice compared to WT. Furthermore, the S1PR1 agonism had no effect on syndecan-1 levels within the time of AMT recovery to normal [7]. We propose that restoring normal AMT in *Apom*^*-/-*^ mice unlikely depends on the recovery of syndecan-1. In addition, syndecan-1 expression is negligible in BECs forming the BBB [35, 71], which may further question its involvement in regulating transcytosis in *Apom*^*-/-*^ mice.

S1PR1 agonism eventually induced slow recovery in syndecan-1 plasma concentrations, but with considerable latency, i.e., at 5 days after recurrent treatment with SEW2871. This seemingly contrasts with the relatively fast, 6-hour recovery of gcx in isolated mesenteric vessels *in vitro* following removal and reinstating S1P in the extracellular environment [53]. In turn, our results agree with other studies reporting gcx restoration within a similar time, i.e., 5-7 days after cytokine-mediated acute degradation in cremaster muscle venules [72]. However, analogies should be made with caution, as the time of gcx recovery likely depends on the tissue preparation (*in vitro, in situ, in vivo*), tissue type (brain, periphery), degree of gcx loss, and the different mechanisms that may underlie the reduction of gcx in acute and chronic dysfunction of the endothelium.

It is plausible that the interplay of S1PR1-gcx-AMT involves other cells supporting BBB function. For instance, S1PR1 is highly expressed in astrocytes [35], and recent evidence suggests that astrocyte presence at the BBB is important to ameliorate gcx loss in disease conditions [73]. However, more research is needed to further relate astrocytes to S1P signaling deficits, gcx reduction and changes in AMT in *Apom*^*-/-*^ mice.

Notably, S1PR1 signaling deficits also lead to an increase in paracellular permeability to small molecules. This has been demonstrated by increased extravasation of sodium fluorescein (∼0.38 KDa) and Alexa488 (∼0.64 kDa) tracers in both *Apom*^*-/-*^ and S1PR1 conditional knock-out mice [7, 74]. Similar to AMT, the paracellular leak could also be mitigated by S1PR1 agonism [7, 74]. However, the mechanism of loss and regain of the BBB paracellular barrier is unlikely tied to the presence of gcx, as the gcx does not constitute the diffusional obstacle for molecules of that size [32].

Regarding methods, the gcx is a fragile structure, seldom preserved *ex vivo* [75-77]. In addition, the gcx structure is highly dynamic, which makes generalizations problematic, as the gcx does not have a persistent and predefined dimension [22, 23]. This is further complicated by BBB exhibiting structural and functional zonation along the microvascular tree [7, 8, 35, 41, 78]. Thus, optimally, both the gcx and the BBB should be studied in the living, intact brain. Here, we applied 2PM imaging in mice, using fluorescent WGA conjugate to study gcx *in vivo*, similar to previous studies [36, 45, 73, 79]. To minimize the influence of background signals, we developed a 2PM photobleaching protocol, but like all analytical approaches, our method also has limitations. They stem from the type of gcx labeling dye used and intrinsic properties of the 2PM imaging technique. Concerning labeling dye, lectin conjugates are low affinity and low specificity markers that bind to disaccharides in gcx GAG chains [25, 44]. Although lectin-based labeling can be applied to study gcx in living cells [22, 23], it does not provide information on the backbone protein component of syndecans and glypicans anchored to the BECs membrane. Thus, decreases of the WGA-conjugate fluorescence reflect only the loss of GAG content rather than the wholesale loss of the gcx. Although complementary methods, such as ELISA, can measure specific components of the gcx in plasma, the readout is not brain-specific and reflects protein levels with contribution from all organs in the body. Thus, new developments in gcx-targeting fluorescent probes are needed to provide a more complete picture of gcx changes in the brain *in vivo*. In addition, WGA-conjugates are not specific to gcx and can label, e.g., microglia and monocytes [80, 81], but this unlikely affected our measurements. Concerning 2PM, the imaged fluorescence signal from gcx is sensitive to the time of WGA-conjugate in the circulation. To minimize the influence of labeling time, we carefully adhered to the timepoints in the experimental timeline, which allowed us to compare the same vessel types between distinct animal groups. However, the imaged signal and bleached fluorescence may also be affected by the depth of the focal plane and the vessel geometry. Therefore, we refrained from quantitative comparisons between distinct vessel types that occupy different cortical depths. Lastly, bleaching phototoxicity might potentially open the BBB; however, we observed no extravasation of blood-circulating WGA-AF594 to the brain. Since we cannot rule out that photobleaching-generated free radicals might affect transcytosis, we avoided the concomitant assessment of the gcx and the AMT.

In summary, our new imaging approach revealed that apoM/S1P signaling deficits cause loss of gcx in cerebral arterioles, with no effect on capillaries and venules. We suggest that although the gcx loss may contribute to the disinhibition of AMT, its presence is not a prerequisite to rescue the impaired BBB with impaired S1P signaling.

## ACKNOWLEDGEMENTS

We would like to acknowledge Micael Lønstrup for his assistance with animal surgery. This study was supported by Lundbeck Foundation Research Initiative on Brain Barriers and Drug Delivery (#R392-2018-2266); Det Frie Forskningsråd (#0602-01965B); Nordea Foundation Grant to the Center for Healthy Aging (#114995); Novo Nordisk Fonden; and Læge Sophus Carl Emil Friis og hustru Olga Doris Friis’ Legat.

